# Assessment of heterotic potential and combining ability of immortal restorer lines derived from an elite rice hybrid, KRH-2, for the development of superior rice hybrids

**DOI:** 10.1101/2020.11.09.373985

**Authors:** Swapnil Ravindra Kulkarni, Balachandran SM, Fiyaz RA, Sruthi K, Divya Balakrishnan, Ulaganathan K, Hari Prasad A.S., Sundaram RM

## Abstract

Present investigation was carried out to assess the heterotic potential and combining ability of immortal restorer lines [consisting of two recombinant inbred lines (RILs) and two doubled haploid lines (DHLs)] developed from an elite rice hybrid, KRH-2 by crossing them with three popular WA-CMS lines, IR58025A, CRMS32A and APMS6A through line × tester analysis. The doubled haploid line 1 (DHL-1) was observed to be a good general combiner for total grain yield per plant (YLD) and other yield component traits and among the CMS lines, IR58025A was observed to be the best combiner as it showed positive significant values for the traits viz., total grain yield per plant, panicle length and spikelet fertility. Higher preponderance of the variance associated with specific combining ability (SCA) as compared to general combining ability (GCA) variance was observed for most of the traits indicated the predominant role of non-additive gene action in the expression of the traits. Out of twelve novel crosses between the immortal restorer lines derived from KRH-2 and the WA-CMS lines, 66.66% (eight crosses) showed significant and desirable SCA effects for the traits viz., total grain yield per plant, days to fifty percent flowering, plant height, flag leaf length, flag leaf width, number of filled grains per panicle and spikelet fertility. Two crosses IR58025A/RIL-24 and CRMS32A/RIL-24 were observed to be the most promising cross combinations showing standard heterosis of >50% for YLD trait (as compared with KRH-2) with higher prevalence of GCA and SCA, respectively. Heterotic yield advantage of IR58025A/RIL-24 and CRMS32A/RIL-24 was 77.05% and 54.74%, respectively over KRH-2 and these can be utilized for developing commercial hybrids. The present study also indicates the potentiality of RILs in providing useful parental lines for developing heterotic hybrids which are hard to get from outside sources in the new intellectual property regime.

## Introduction

Rice (*Oryza sativa* L.) is one of the world’s largest food crop and Asian countries are predominantly dependent on rice for their nutritional and calorific needs.It was predicted by [1] that by 2030, the demand for rice will escalate to 852 million tons. Moreover, the rice breeders have to face many challenges in increasing the rice production due to declining land area for rice cultivation, shortage of labor and water and imminent threats posed by biotic and abiotic stresses [2]. Enhancement of rice productivity through innovative genetic approaches such as hybrid rice technology offers a hope of lessening the gap between rice demand and its production [3]. Hybrid rice breeding aims to break the yield barrier by exploiting the phenomenon of heterosis in order to increase the yield potential beyond the level of high yielding varieties. Being a staple food crop of India, rice is grown in about 44.15 million hectares with annual production of 116.47 million tonnes and productivity of 2638 Kg ha^-1^[4].A total of 107 hybrids have been released for commercial cultivation in India [5] but hybrid rice occupies only 3 million hectares of the total rice in the country [6]. Though these rice hybrids yielded 15-20% more than the semi-dwarf inbred varieties [7, 8], their adoption is not up the expected level due to various reasons. Unattractive yield advantage of hybrids, issues related to quality, non-diverse sources of restorers and CMS lines, lack of favorable policies, pricing discrimination against hybrids are some of the reasons for lower adoption of hybrids in India. [9]identified the lack of genetic diversity in the existing rice gene pool and lower to moderate yield advantages in novel hybrids [10] as some of the prime reasons for lesser adoption of hybrid rice for cultivation. As opined by [11], it is possible to overcome the existing yield plateau by broadening the genetic base of rice gene pool through identification of efficient-diverse restorer (R) lines and stable CMS (A) lines with higher out crossing ability and by envisaging novel crosses between them using the three line hybrid system popularly known as cytoplasmic genic male sterile (CGMS) line system. Traditionally, the production of hybrid rice has been based on CMS i.e., the A line, maintainer (B) line and restorer (R) line. This involved the crosses between CMS lines and R lines which produced novel fertile F_1_ offsprings. In hybrid rice breeding, there is need to develop an array of diverse restorers as the available CMS lines will suffice to make several crosses. It was noted that the fertility restoration in hybrids was as a consequence of dominant fertility restorer genes (*Rf* genes) from R lines [12-16]. Plant breeders face enormous challenge in the development of heterotic hybrids due to lower diversity among parental lines. Promising parental lines have to be identified based on their combining ability and superior hybrids through their *per se* performance. Therefore, through line × tester (L×T) mating design, the potential of individual parental line to transmit the genetic information to novel offsprings can be quantified [17] enabling the assessment of general combining ability (GCA) and specific combining ability (SCA) [3, 18]. GCA prominently associated with additive gene action, evaluates mean performance of a line in various crosses and identifies superior parental lines on the basis of mean value. This aids in the identification of promising parental lines and there by producing heterotic hybrids [3]. The SCA on the other hand, is the result of non-additive in gene action and manifests essentially from over dominance, dominance and epistatic effects [19]. It primarily helps in the identification of promising hybrids by assessing the positive or negative genetic value of the expected mean performance of the parental crosses [20, 21]. The present study aimed at identification of potential restorers from DHL and RIL population derived from an elite rice hybrid KRH-2, assessing *per se* performance of novel hybrids for yield and its component traits, understanding the genetic basis of hybrid improvement through GCA-SCA studies and identification of promising novel heterotic hybrids for commercialization.

## Materials and methods

### Experimental material

A popular rice hybrid, Karnataka Rice Hybrid-2 (KRH-2) with following characteristics i.e., medium duration, long-bold grain type, suitable for irrigated ecology and developed by Zonal Agricultural Research Station (ZARS), Mandya, Karnataka, India and with a high yield potential along with its parents, IR58025A (in this study we have used IR58025B line for agro-morphological evaluation as IR58025A does not set seeds) and KMR-3R, was used as the experimental material.

### Development of KRH-2 doubled haploid lines (DHLs) and recombinant inbred lines (RILs) population and their agro-morphological evaluation

Genetically pure seeds of parents (IR58025A and KMR-3R) and the elite rice hybrid KRH-2 were grown in the research farm of ICAR-Indian Institute of Rice Research (ICAR-IIRR), Hyderabad India, during the dry season of 2015, sowing on well-puddled, leveled and raised nursery beds, 15-20 days old seedlings were transplanted in field with a spacing of 15×20cm (6 rows×20 hills) and the recommended package of practices were adopted for growing the plants. A total of 125 regenerated true and highly stable doubled haploid lines (DHLs) (D_0_) were developed following the standardized protocol of [22]. Similarly, for the development of RIL population, the F_1_ hybrid seeds (produced from the crosses between IR58025A and KMR-3R)were self-pollinated to produce F_2_ progenies. The F_2_ progenies were later advanced to further generations in the field through single seed descent (SSD) method in subsequent seasons [23]. A total of 105 individuals consisted of RIL population. Both the DHL and RIL populations were grown for three consecutive seasons (dry season 2016, D_2_ generation of DHL population and F_3_ generation of RIL population; wet season 2016, D_3_ generation and F_4_ generation; dry season 2017, D_4_ generation and F_5_ generation) for analyzing their genetic variability with respect to key agro-morphological traits viz., days to fifty percent flowering (DFF), total grain yield per plant (YLD), total number of grains per panicle (GP), filled grains per panicle (FGP), test (1000 grains) weight (TGW), panicle weight (PW), plant height (PH), panicle length (PL), flag leaf length (FLL), flag leaf width (FLW), number of productive tillers (NPT) and biomass (BM). Data was recorded from five healthy plants of middle row of each line, as per the standard evaluation system recommended by [24].

### Selection of restorers among KRH-2 derived DHL and RIL population and their fertility restoration assessment

A total of 40 lines (among both the immortal populations) were selected based on the presence of fertility restoration genes and performance of the lines for various traits. Of the total 40 lines, 16 DHLs (D_5_) which are high-yielding and semi-tall (plant height of 110±5 cm) to tall (plant height of 125±5 cm) and 24 RILs (F_6_) i.e., 12 high and 12 low yielding, semi-tall were selected. Molecular screening of selected RILs and DHLs for the presence of fertility restoration genes using functional markers namely RMS-SF21-5 and RMS-PPR9-1, specific for major fertility restorer genes, *Rf3* locus and *Rf4* locus, respectively, was undertaken as described in [25]. For this genomic DNA was extracted from IR58025A, IR58025B, KMR-3R along with the selected RILs and DHLs as described in [26] and allelic status was observed. Further, the allelic status of these lines in terms of their amplification for *Rf3-Rf4* loci was correlated with fertility restoration potential in novel hybrids derived from them [25]. The lines were considered as complete restorers if their restoration potential in novel hybrids was observed to be greater than or equal to 70% [15]. The selected lines were test crossed with the WA-CMS line, IR58025A during the wet season of 2017-18 using line×tester mating design [18]. The novel hybrids produced from these crosses along with their parents (D_6_ and F_7_ generations), standard varietal checks namely Akshayadhan (AKD) and Varadhan (VRD) were assessed for their agro-morphological trait performance and heterosis studies in dry season of 2018-2019. Novel hybrids produced from two high yielding DHLs namely DHL-1 and DHL-2 and those produced from highest yielding RIL-1 and low yielding RIL-24, were observed to be heterotic than KRH-2 and these four restorers were further selected for test crossing with other CMS lines.

### Test crossing of selected restorers with popular WA-CMS lines

In wet season of 2018, four selected parental lines viz., DHL-1 (D_7_), DHL-2 (D_7_), RIL-1 (F_8_) and RIL-24 (F_8_) were test crossed with three popular CMS lines viz., IR58025A, CRMS32A and APMS6A to assess the general combining ability (GCA) of parents and specific combining ability (SCA) of crosses using line × tester mating design [18].

### Assessment of general combining ability (GCA) of parents and specific combining ability (SCA) of crosses and estimation of heterosis of the newly derived hybrids

In dry season of 2019-2020, 12 novel hybrids along with their parents including IR58025B, CRMS32B and APMS6B (Maintainers of CMS lines IR58025A, CRMS32A and APMS6A); four restorer lines DHL-1 (D_8_), DHL-2 (D_8_), RIL-1 (F_9_), RIL-24 (F_9_) and five hybrid checks, viz., KRH-2, US312, US314, PA6444, HRI174 were evaluated for combining ability and heterosis studies [27]. Following the randomized complete block design (RCBD) in the experimental field, the data was recorded from ten healthy plants of middle row of each line from two replications (five plants from each replication), as per the standard evaluation system (IRRI, 2002) on 12 important yield attributing traits viz., days to fifty percent flowering (DFF); plant height (PH); flag leaf length (FLL); flag leaf width (FLW); number of productive tillers (NPT); panicle length (PL); total number of grains per panicle (TNGPP); number of filled grains per panicle (NFGPP); number of unfilled grains per panicle (NUFGPP); spikelet fertility in percentage (SFP); test (1,000 grains) weight (TGW) and total grain yield per plant (YLD). Estimates of GCA and SCA [28] were computed using R language statistical software version Ri386 3.3.2.

### Statistical analysis

Data of twelve agro-morphological and yield related traits were subjected to statistical analysis viz., Analysis of variance (ANOVA), genotypic and phenotypic correlations, genetic variability estimates were analyzed using SAS version 9.2 (SAS Institute Inc., Cary, NC, USA). L×T mating design ANOVA, genetic variances and heritability estimates were computed using R language statistical software version Ri386 3.3.2. Mean, standard deviation (SD), standard error (SE) and coefficient of variation in percentage (CV %) were computed using Microsoft Excel package. Heterobeltiosis, mid-parent heterosis [29] and standard heterosis were estimated using standard formulae [30, 31].

## Results

### Selection of parents based on assessment of the novel hybrids

Based on the assessment of yield heterosis data (evaluated in dry season of 2018-2019; S1 Table and S2 Table) of the novel hybrids, it was observed that two crosses namely IR58025A/DHL-1 and IR58025A/DHL-2 demonstrated very high positive standard heterosis for total grain yield per plant (YLD) as compared to varietal checks (Akshayadhan [AKD] andVaradhan[VRD]) and standard hybrid check(KRH-2). The standard YLD heterosis of the hybrid IR58025A/DHL-1 was observed to be 63.04, 48.61 and 25.21% over the checks AKD, VRD and KRH-2 respectively. While, the hybrid IR58025A/DHL-2 recorded 47.65, 34.58 and 10.20% over the checks AKD, VRD and KRH-2 respectively.Further, both DHL-1 and DHL-2, showed the presence of both *Rf3* and *Rf4* (the major fertility restoration genes), when analyzed with functional markers specific for these genes. This was corroborated with the fertility restoration data of both these lines, wherein high values of spikelet fertility were observed in the derived hybrids (viz., 85.42% for DHL-1 and 81.36% for DHL-2). The hybrids derived from the cross IR58025A and the highest yielding RIL-1, demonstrated a yield advantage of 177.34% over AKD, 162.23% over VRD and 111.38% over KRH-2. Another hybrid derived from the cross IR58025A and one of the low yielding RILs, viz., RIL-24 registered a yield advantage of 38.35% over AKD, 46.32% over VRD and 11.52% over KRH-2. Analysis of fertility restoration of these two lines with respect to*Rf3-Rf4* specific markers revealed the presence of both the loci in them and the results corresponded to the spikelet fertility values of the hybrids derived from the two RILs (80.97% and 85.99% with respect to RIL-1 and RIL-24, respectively).

### Agro-morphological performance of novel hybrids

Novel hybrids that were developed between three popular CMS lines (IR58025A, CRMS32A and APMS6A) and four selected restorers (DHL-1, DHL-2, RIL-1, RIL-24) were evaluated for their agro-morphological performance in the dry season of 2019-2020 and the details of which are described below.

### Genetic variability estimates

As shown in S3 Table, in the novel hybrids, high phenotypic (Vp) and high genotypic variance (Vg) was observed for two traits, viz., number of fertile grains per panicle (NFGPP, Vp = 1603.84 and Vg = 1447.75) and total number of grains per panicle (TNGPP, Vp = 1556.64 and Vg = 1343.41). The trait flag leaf width (FLW, Vp = 0.04 and Vg = 0.02) was observed to have the lowest phenotypic-genotypic variance. Collectively for all the traits, the phenotypic variance was observed to be higher than the genotypic variance. Based on GCV-PCV values, the traits were classified into three categories, viz., low, moderate and high. Higher (>20%) genotypic coefficient of variance (GCV)-phenotypic coefficient of variance (PCV) values was observed for the traits, NPT, TNGPP, NFGPP, NUFGPP and YLD. Moderate (10%-20%) of GCV-PCV values were observed for the traits DFF, PH, FLL, FLW and SFP while low (0%-10%) of GCV-PCV values were observed for traits viz., PL and TGW. All the traits under study recorded high broad sense heritability (*H*^*2*^) values except FLW. Genetic advance (GA) was observed within the range of 0.25 (FLW) to 74.46 (NFGPP). High GAM values were observed for traits viz., DFF, PH, FLL, NPT, TNGPP, NFGPP, SFP and YLD. High *H*^*2*^and high GAM values were observed for the traits DFF, PH, FLL, NPT, TNGPP, NFGPP, SFP and YLD. Moderate *H*^*2*^ with moderate GAM was observed for FLW trait and high *H*^*2*^ with moderate GAM was observed for PL trait.

### Correlation between different traits

Positive correlation was observed between YLD and most of its component traits (S4 Table). Significant positive correlation was observed between YLD and NPT (r = 0.23), FLL (r = 0.05), PL (r = 0.38), TNGPP (r = 0.14), SFP (r = 0.45) and TGW (0.24) at 1% level of significance, while a positive correlation was observed between YLD and FLW (r = 0.10), NFGPP (r = 0.28) at 5% level of significance. Negative correlation was noticed between YLD and DFF (r = -0.34) and NUFGPP (r = -0.31) at 1% and 5% levels of significance, respectively. Significant correlation was observed between the traits TNGPP and NPT (r = 0.26) and PL (r = 0.42) at 1% and 5%, levels of significance, respectively. A strong positive correlation between SFP and TNGPP (r = 0.17) and negative correlation between NUFGPP (r = -0.78) was observed at 1% level of significance, while significant positive correlation was observed between TGW and TNGPP (r = 0.51) at 1%, NPT (r = 0.26), PL (r = 0.09), NFGPP (r = 0.19) and SFP (r = 0.10) at 5% level of significance.

### Estimation of genetic variances and heritability in novel crosses

Genetic parameters such as variance due to general combining ability (GCA), specific combining ability (SCA), GCA variance ratio, additive genetic variance, dominance genetic variance, degree of dominance and broad sense-narrow-sense heritability were estimated among the hybrids. It was observed that for all traits under study, the variance due to SCA (σ^2^sca) was higher than GCA (σ^2^gca). Also, the dominance genetic variance (δ^2^D) was larger than the additive genetic variance (δ^2^A) for all the traits. These results are supported with GCA variance ratio (σ^2^gca / σ^2^sca) being less than 1 for all traits and degree of dominance [(δ^2^D/ δ^2^A)^1/2^] being less than 1 for trait viz., TNGPP, NUFGPP, PH, YLD, DFF, FLL and for remaining traits viz., FLW, NPT, PL, NFGPP, SFP and TGW the value was greater than 1. Broad sense heritability (*H*^*2*^ (%)) was observed in the range of 56.16% (FLW) to 99.78% (DFF) whereas the narrow sense heritability (*h*^*2*^ (%)) was in the range 14.53% (PL) to 49.10% (DFF) (S5 Table).

### Estimation of heterosis among the novel hybrids

Test crosses carried out between three popular, commonly deployed WA-CMS lines, viz., IR58025A, CRMS32A, APMS6A and the four selected restorers, viz., RIL-1, RIL-24, DHL-1 and DHL-2 produced twelve novel hybrids, which were evaluated in wet season of 2018. In the present study, variance analysis for all the characters revealed significant variation among the genotypes studied (S6 Table).The coefficient of variation (CV %) values were observed to be <20% for most of the traits except NPT, TNGPP, NFGPP and YLD. The agro-morphological performance of the novel hybrids, parents along with standard hybrid checks is presented in Table 1. The mean values for eleven yield component traitsin parents, standard hybrid checks and in novel hybrids were as follows: DFF: 96 (range: 81-122), PH: 99.86 cm (range: 73.98-124.00 cm), FLL: 28.64 cm (range: 20.87-43.01 cm), FLW: 1.37 cm (range: 1.10-1.97 cm), NPT: 17 (range: 15-18), PL: 22.89 cm (range: 19-28.01 cm), TNGPP: 120 (range: 67-211), NFGPP: 83 (range: 39-173), NUFGPP: 30 (range: 6-81), SFP%: 73.76% (range: 45.90%-93.88%), TGW: 20.17 g (range: 12.37-23.20 g) and YLD: 33.20 g (16.17-50.23 g).

**Table 1:** Agro-morphological data of novel hybrids derived from the test cross between selected DHL-RIL restorers with three popular WA-CMS lines along with parents and checks

| Material/Cross description | Generation | DFF±SE | PH±SE | FLL±SE | FLW±SE | NPT±SE | PL±SE | TNGPP±SE | NFGPP±SE | NUFGPP±SE | SFP±SE | TGW±SE | YLD±SE |
| --- | --- | --- | --- | --- | --- | --- | --- | --- | --- | --- | --- | --- | --- |
| APMS6A/RIL-1 | Novel hybrid (F<sub>1</sub>) | 107±0.00 | 121.37±0.05 | 36.35±0.33 | 1.19±0.44 | 15±0.52 | 25.40±0.22 | 126±0.36 | 64±0.66 | 62±0.25 | 50.98±0.33 | 21.02±0.28 | 20.10±0.48 |
| APMS6A/RIL-24 |  | 89±0.00 | 103.82±0.25 | 29.60±0.47 | 1.48±0.92 | 16±0.23 | 23.15±0.48 | 123±0.56 | 90±0.35 | 33±0.22 | 73.38±0.23 | 20.28±0.63 | 29.20±0.36 |
| APMS6A/DHL-2 |  | 91±0.00 | 73.98±0.45 | 43.01±0.41 | 1.13±0.42 | 18±0.22 | 19.01±0.22 | 67±0.58 | 40±0.21 | 27±0.36 | 59.84±0.54 | 22.46±0.44 | 16.17±0.22 |
| APMS6A/DHL-1 |  | 84±0.00 | 77.52±0.33 | 24.00±0.35 | 1.31±0.66 | 18±0.42 | 20.33±0.41 | 132±0.74 | 119±0.36 | 13±0.21 | 90.16±0.63 | 19.66±0.32 | 42.21±0.63 |
| IR58025A/RIL-1 |  | 111±0.00 | 97.37±0.68 | 28.54±0.68 | 1.36±0.42 | 16±0.47 | 26.48±0.85 | 116±0.36 | 95±0.84 | 21±0.52 | 81.84±0.45 | 18.87±0.45 | 28.68±0.33 |
| IR58025A/RIL-24 |  | 122±0.00 | 97.16±0.74 | 32.37±0.47 | 1.34±0.42 | 15±0.36 | 28.01±0.36 | 211±0.44 | 173±0.74 | 38±0.36 | 81.33±0.85 | 19.36±0.33 | 50.23±0.21 |
| IR58025A/DHL-2 |  | 81±0.00 | 92.00±0.55 | 20.87±0.66 | 1.20±0.36 | 16±0.21 | 23.67±0.75 | 99±0.21 | 87±0.51 | 12±0.21 | 87.60±0.66 | 23.20±0.77 | 32.29±0.63 |
| IR58025A/DHL-1 |  | 107±0.00 | 95.83±0.64 | 32.03±0.12 | 1.61±0.32 | 16±0.33 | 24.73±0.35 | 109±0.36 | 84±0.33 | 25±0.85 | 76.66±0.21 | 20.75±0.98 | 27.88±0.55 |
| CRMS32A/RIL-1 |  | 96±0.00 | 86.77±0.41 | 29.44±0.47 | 1.10±0.45 | 16±0.54 | 23.86±0.88 | 103±0.88 | 64±0.14 | 39±0.66 | 62.32±0.96 | 20.51±0.66 | 21.00±0.45 |
| CRMS32A/RIL-24 |  | 102±0.00 | 108±0.77 | 27.33±0.36 | 1.43±0.42 | 18±0.36 | 23.33±0.91 | 141±0.96 | 118±0.08 | 23±0.54 | 83.40±0.66 | 20.67±0.23 | 43.90±0.36 |
| CRMS32A/DHL-2 |  | 106±0.00 | 90.26±0.45 | 28.33±0.11 | 1.40±0.44 | 16±0.33 | 23.30±0.45 | 139±0.31 | 109±0.55 | 30±0.36 | 78.54±0.36 | 22.92±0.33 | 40.93±0.21 |
| CRMS32A/DHL-1 |  | 97±0.00 | 101.33±0.36 | 29.33±0.32 | 1.70±0.43 | 16±0.47 | 21.00±0.36 | 189±0.22 | 168±0.36 | 21±0.33 | 88.99±0.45 | 15.23±0.21 | 39.97±0.33 |
| APMS6B | Maintainer | 94±0.00 | 105.67±0.84 | 25.67±0.49 | 1.50±0.44 | 13±0.36 | 20.67±0.24 | 156±0.20 | 106±0.21 | 50±0.84 | 66.83±0.63 | 16.73±0.32 | 16.73±0.21 |
| IR58025B | Maintainer | 88±0.00 | 85.00±0.33 | 26.00±0.52 | 1.13±0.53 | 17±0.44 | 21.83±0.47 | 84±0.19 | 39±0.25 | 45±0.78 | 45.90±0.32 | 19.40±0.44 | 12.86±0.95 |
| CRMS32B | Maintainer | 92±0.00 | 87.93±0.48 | 26.60±0.34 | 1.17±0.85 | 14±0.47 | 21.00±0.36 | 112±0.88 | 75±0.77 | 37±0.66 | 67.36±0.45 | 18.06±0.96 | 18.96±0.36 |
| RIL-1 | Tester (F<sub>7</sub>) | 88±0.00 | 124.00±0.96 | 24.00±0.56 | 1.23±0.43 | 18±0.23 | 22.67±0.55 | 94±0.71 | 88±0.44 | 6±0.95 | 93.88±0.99 | 22.57±0.32 | 35.64±0.66 |
| RIL-24 | Tester (F<sub>7</sub>) | 103±0.00 | 128.67±0.44 | 29.00±0.44 | 1.43±0.96 | 14±0.47 | 23.67±0.36 | 88±0.36 | 73±0.95 | 15±0.66 | 82.52±0.69 | 19.77±0.11 | 20.20±0.32 |
| DHL-2 | Tester (D<sub>4</sub>) | 91±0.00 | 110.67±0.42 | 34.00±0.25 | 1.97±0.88 | 13±0.69 | 25.33±0.44 | 189±0.44 | 108±0.35 | 81±0.35 | 57.43±0.32 | 18.73±0.21 | 16.77±0.02 |
| DHL-1 | Tester (D<sub>4</sub>) | 85±0.00 | 94.33±0.36 | 25.33±0.54 | 1.37±0.43 | 14±0.35 | 21.33±0.96 | 91±0.36 | 85±0.24 | 6±0.45 | 93.31±0.65 | 21.87±0.32 | 25.90±0.22 |
| HRI174 | Hybrid check 1 | 97±0.00 | 100.00±0.45 | 29.00±0.77 | 1.50±0.44 | 10±0.44 | 20.00±0.87 | 98±0.21 | 69±0.32 | 29±0.36 | 69.84±0.75 | 18.53±0.33 | 25.60±0.47 |
| PA6444 | Hybrid check 2 | 97±0.00 | 103.67±0.48 | 28.00±0.36 | 1.23±0.21 | 15±0.87 | 22.33±0.96 | 75±0.69 | 52±0.33 | 23±0.84 | 68.91±0.36 | 20.59±0.56 | 19.68±0.65 |
| US314 | Hybrid check 3 | 87±0.00 | 92.00±0.37 | 27.33±0.42 | 1.37±0.31 | 14±0.56 | 20.83±0.55 | 80±0.22 | 45±0.25 | 35±0.36 | 56.91±0.44 | 20.47±0.96 | 17.38±0.55 |
| US312 | Hybrid check 4 | 90±0.02 | 101.33±0.48 | 23.00±0.66 | 1.53±0.24 | 14±0.41 | 25.67±0.36 | 140±0.47 | 100±0.66 | 40±0.22 | 71.39±0.36 | 20.13±0.33 | 25.77±0.36 |
| KRH-2 | Hybrid check 5 | 97±0.00 | 118.00±0.71 | 29.00±0.45 | 1.13±0.63 | 13±0.36 | 22.33±0.31 | 128±0.66 | 104±0.21 | 24±0.32 | 80.97±0.84 | 21.20±0.32 | 28.73±0.55 |
|  | Mean | 97 | 99.05 | 28.91 | 1.40 | 15 | 23.23 | 137 | 107 | 30 | 75.64 | 20.26 | 27.08 |
|  | SD | 10.25 | 13.12 | 4.53 | 0.22 | 2.20 | 2.51 | 70.01 | 69.86 | 17.25 | 13.13 | 2.04 | 9.84 |
|  | SE | 2.05 | 2.62 | 0.91 | 0.04 | 0.44 | 0.50 | 14.00 | 13.97 | 3.45 | 2.63 | 0.41 | 1.97 |
|  | CV% | 10.57 | 13.25 | 15.67 | 15.36 | 14.28 | 10.82 | 51.07 | 65.49 | 56.80 | 17.36 | 10.05 | 36.32 |

**276** DFF-Days to fifty percent flowering (in days); PH-Plant Height (cm), FLL-Flag Leaf Length (cm); FLW-Flag Leaf width (cm); NPT-Number of Productive Tillers; PL-Panicle Length (cm); TNGPP- **277** Total Number of Grains Per Panicle; NFGPP-Number of Filled Grains per Panicle; NUFGPP-Number of Unfilled Grains per Panicle; SFP-Spikelet Fertility in Percentage; TGW-Test (1,000) Grain Weight **278** (g); YLD-Total grain Yield per plant (g); SD-Standard deviation; SE-Standard error; CV%-Coefficient of variation in percentage

Difference in the magnitude of three heterosis categories namely, mid-parent heterosis (MPH), better parent heterosis (BPH) and standard heterosis (SH) for YLD and its allied components was calculated at 1% and 5% level of significance. The average YLD of novel hybrids (32.71 g) was higher than that of the average YLD of the isogenic counterpart of CMS lines (i.e., maintainer lines) (17.56 g), of restorers (average YLD-24.02 g) and that of the standard hybrid checks (average YLD-20.68 g). For YLD trait, seven crosses viz., IR58025A/RIL-24, CRMS32A/RIL-24, APMS6A/RIL-24, CRMS32A/DHL-1, APMS6A/DHL-1, IR58025A/DHL-2 and CRMS32A/DHL-1 were observed to be positively heterotic as compared to the standard hybrid checks, mid and better parents. The values for standard heterosis ranged from 189.01% (IR58025A/RIL-24 vis-à-vis US314) to 1.09% (IR58025A/RIL-1 vis-à-vis KRH-2) (S7 Table). The details of heterosis of the cross combinations with respect to YLD trait is presented in S7 Table.

For NPT trait, novel hybrids from all crosses were observed to have more number of productive tillers than all parents and checks. The magnitude of heterosis was observed in the range of -16.67% (BPH) to 80% (HRI174). Range of NPT in these hybrids was 15-18. Out of twelve novel hybrids, two hybrids derived from the crosses IR58025A/RIL-24 and CRMS32A/DHL-1 were positively heterotic for all the categories of TNGPP heterosis. The trait improvement in these hybrids was observed in the range of 40.58% (MPH) to 170.51% (SH over PA6444). Test weight of three hybrids namely IR58025A/DHL-2 (23.20 g), CRMS32A/DHL-2 (22.92 g) and APMS6A/DHL-2 (22.46 g) surpassed all parents and checks whose heterosis was in the range of 9.99% (SH over PA6444) to 31.87% (MPH). Heterotic details of all cross combinations for the traits other than YLD are presented in S8 Table.

Early flowering and high yielding hybrid is a desirable combination and in this study four such novel hybrids were identified. IR58025A/DHL-2, a high-yielding (32.29 g) and was an early flowering hybrid (DFF, 81 days) than US314 (early duration hybrid); CRMS32A/DHL-1 with YLD of 39.97 g had DFF (97 days) similar to PA6444 and HRI174 (medium duration hybrids) but was late than KRH-2; CRMS32A/RIL-24 (YLD, 43.90 g; DFF, 102 days) was early flowering than the better parent (CRMS32A) but was late than medium duration hybrids (KRH-2, PA6444, HRI174); APMS6A/RIL-24 (YLD, 29.20 g; DFF-89 days) was early flowering than medium duration hybrids (KRH-2, PA6444, HRI174) (Table 1).

### Combining Ability analysis

The Analysis of Variance of the parental lines and checks using L×T mating design is presented in Table 2. The variances due to parents were significant for the following traits: DFF, PH, FLW, NPT, PL, NFGPP, NUFGPP, TNGPP, TGW, SFP at *P*<0.001 and FLL, YLD at *P*<0.05. Variances due to parents versus crosses were observed to be significant (*P*<0.001) for traits namely DFF, PH, FLL, NPT and YLD. Variances due to crosses were significant (*P*<0.001) for all traits except SFP. The L × T effect was significant (*P*<0.001) for traits DFF, PH, FLL, NPT and TNGPP whereas the effect was significant (*P*<0.01) for traits NFGPP, NUFGPP and TGW. Among the proportion of contribution of lines and testers, it was observed that the lines had contributed (>40%) for traits namely NPT, PL, SFP whereas the testers had contributed for traits namely FLW and TGW. A major proportion of trait contribution was observed for traits namely DFF, PH, FLL, NFGPP, TNGPP, TGW and YLD by L×T.

**Table 2:** Analysis of variance (ANOVA) for yield and its associated traits

| Source | df | DFD | PH | FLL | FLW | NPT | PL | NFGPP | NUFGPP | TNGPP | SFP | TGW | YLD |
| --- | --- | --- | --- | --- | --- | --- | --- | --- | --- | --- | --- | --- | --- |
| Replication | 2 | 0 | 0.06 | 0.3 | 0.54 | 0.04 | 0.17 | 0.38 | 0.5 | 0.32 | 1.13 | 0.49 | 2.43 |
| Genotype | 19 | 59.50*** | 21.38*** | 6.98*** | 6.31*** | 26.97*** | 4.05*** | 5.26*** | 11.12*** | 5.35*** | 7.24*** | 5.48*** | 13.13*** |
| Parents | 7 | 16.83*** | 25.33*** | 2.87* | 9.72*** | 8.22*** | 2 | 2.32 | 21.23*** | 5.28*** | 11.60*** | 5.46*** | 3.55* |
| Parents vs. crosses | 1 | 153.99*** | 45.45*** | 13.70*** | 0.34 | 160.66*** | 7.68* | 7.14* | 4.59 | 2.6 | 5.07 | 2.05 | 125.28*** |
| Crosses | 11 | 78.06*** | 16.68*** | 8.98*** | 4.68*** | 26.70*** | 5.02*** | 6.96*** | 5.27*** | 5.64*** | 4.65** | 5.80*** | 9.03*** |
| Lines | 2 | 1.06 | 0.03 | 0.62 | 0.84 | 12.26 | 7.21 | 2.44 | 0.44 | 1.22 | 6.08 | 0.34 | 0.67 |
| Testers | 3 | 0.73 | 1.08 | 0.1 | 3.21 | 0.4 | 4.5 | 1.02 | 1.27 | 0.66 | 2.23 | 2.39 | 0.24 |
| Line × testers | 6 | 83.14*** | 19.72*** | 13.04*** | 2.97* | 9.25*** | 1.63 | 5.48** | 5.41** | 5.95*** | 2.05 | 4.60** | 12.28 |
| Error | 50 | ----- | ----- | ----- | ----- | ----- | ----- | ----- | ----- | ----- | ----- | ----- | ----- |
| Contribution of lines (%) |  | 20.62 | 0.64 | 16.46 | 9.77 | 77.26 | 42.52 | 35.06 | 8.35 | 23.45 | 48.92 | 4.9 | 16.76 |
| Contribution of testers (%) |  | 21.28 | 34.86 | 4.28 | 55.62 | 3.83 | 39.79 | 21.97 | 35.59 | 18.99 | 26.96 | 51.83 | 9.11 |
| Contribution of L × T(%) | | 58.09 | 64.48 | 79.24 | 34.59 | 18.89 | 17.67 | 42.96 | 56.04 | 57.55 | 24.1 | 43.25 | 74.12 |

### General Combining Ability (GCA) analysis

A brief account of GCA effects of all CMS lines and restorers for various traits namely YLD, NPT, TNGPP and TGW are presented (Table 3). Neither the lines nor the testers could demonstrate positive GCA collectively for all the traits. For total grain yield per plant (YLD) trait, CMS line IR58025A was observed to be the best combiner and DHL-1 (at *P*< 0.001)was the best combiner among the testers. For trait number of productive tillers (NPT), APMS6A was observed to be the best combiner with GCA value of 6.22 (*P*< 0.001) followed by DHL-2 (1.67, *P*< 0.01). For TNGPP trait, highest GCA effect was recorded with RIL-24 (GCA value: 21.43, *P*< 0.01) followed by CRMS32A (GCA value: 16.2, *P*< 0.05). For one of the most crucial YLD contributing trait i.e., TGW, the following observation was made: DHL-2 was the best combiner (GCA value: 2.10, *P*< 0.001) and APMS6A, the best combiner among the lines (0.55, non-significant).

**Table 3:** Estimation of general combining ability effects of parents for yield and its associated traits

| Parents | DFF | PH | FLL | FLW | NPT | PL | NFGPP | NUFGPP | TNGPP | SFP | TGW | YLD |
| --- | --- | --- | --- | --- | --- | --- | --- | --- | --- | --- | --- | --- |
| IR58025A | 5.91*** | 0.13 | -1.58 | 0.02 | -5.28*** | 2.11*** | 12.97 | -3.31 | 9.65 | 6.16*** | -0.09 | 6.13*** |
| CRMS32A | 0.91*** | 1.14 | -1.42 | 0.05 | -0.93 | -0.60 | 17.76*** | -1.57 | 16.2* | 3.96 | -0.46 | -1.45*** |
| APMS6A | -6.83*** | -1.27 | 3.01** | -0.07 | 6.22*** | -1.50** | -30.75* | 4.89 | -25.85* | -10.0*** | 0.55 | -4.64* |
| S.E (line) | 0.68 | 1.58 | 0.91 | 0.04 | 0.54 | 0.54 | 7.2 | 2.77 | 8.38 | 2.48 | 0.41 | 1.91 |
| RIL-1 | 5.33*** | 6.38*** | 1.4 | -0.13 | -0.38 | 1.76*** | -18.5*** | 11.34** | -7.17 | -9.11*** | -0.16 | 1.66 |
| DHL-1 | -3.33*** | -3.89*** | -1.58 | 0.18*** | -1.23* | -1.45** | 14.18* | -5.6* | 8.57 | 4.21*** | -1.74*** | 4.61*** |
| DHL-2 | -6.80*** | -10.0*** | 0.42 | -0.11* | 1.67** | -1.48** | -15.73** | -7.1 | -22.84** | 1.76 | 2.10*** | -3.61*** |
| RIL-24 | 5.08*** | 7.54*** | -0.24 | 0.06 | -0.05 | 1.17* | 20.07** | 1.36 | 21.43** | 3.13 | -0.19(6) | -2.66*** |
| S. E. (tester) | 0.78 | 1.82 | 1.05 | 0.04 | 0.63 | 0.63 | 8.4 | 3.2 | 9.68 | 2.86 | 0.47 | 2.2 |

DFF-Days to fifty percent flowering; PH-plant height (cm), FLL-Flag leaf length (cm); FLW-Flag leaf width (cm); NPT-Number of productive tillers; PL-panicle length (cm); NFGPP- Number of filled grains per panicle; NUFGPP-Number of unfilled grains per panicle TNGPP- Total number of grains per panicle; SFP-Spikelet fertility in percentage; TGW-test (1,000) grain weight (g); YLD-Total grain yield per plant (g), S.E.-Standard Error; \* $P < 0.05$ ; \*\* $P < 0.01$ ; \*\*\* $P < 0.001$ , The significant GCA values (indicated by \*) for each trait is presented.

### Specific Combining Ability (SCA) analysis

Similar to the results obtained in GCA analysis, none of the novel hybrid combination could demonstrate a positive SCA effect for all traits under study (Table 4). For the traits NPT, TGW and YLD, the highest SCA values of 3.68, 2.29 and 14.15 (*P*< 0.001), respectively, was observed between to the hybrid derived from the cross, IR58025A and DHL-1. For all cross combinations, the range of SCA values for NPT trait was observed to be -3.76 to 3.68 (*P*< 0.01) and 58.33% (i.e., seven cross combinations) of hybrids demonstrated positive SCA effect for this trait. For the trait TNGPP, the range of SCA values was observed to be between -36.53 and 50.84. It was noted that, 41.66% of crosses (i.e., five crosses) demonstrated positive SCA effect among which, the hybrid derived from IR58025A/RIL-24 recorded the highest SCA value of 50.84 (*P*< 0.01). For TGW trait, apart from the hybrid derived from IR58025A/DHL-1 observed to have the highest SCA effect (2.29, *P*< 0.01), 41.66% (i.e., five) of other crosses demonstrated a positive SCA value. Lastly for YLD trait, apart from the cross combination IR58025A/DHL-1 (with highest SCA effect value, 14.15, *P*< 0.001), 33.33% (i.e., four) of remaining crosses showed a positive SCA value. The range of SCA values of all crosses was in the range of -14.3 to 14.15 (*P*< 0.001). SCA effect of all crosses for remaining traits is presented in the Table 4.

**Table 4:** Estimation of specific combining ability effects of various crosses for yield and its associated traits

| Entry | DFF | PH | FLL | FLW | NPT | PL | NFGPP | NUFGPP | TNGPP | SFP | TGW | YLD |
| --- | --- | --- | --- | --- | --- | --- | --- | --- | --- | --- | --- | --- |
| IR58025A/DHL-1 | 5.08*** | 4.12* | 5.16** | 0.04 | 3.68** | 0.59 | -38.3*** | 6.77 | -31.57** | -9.28*** | 2.29** | 14.15*** |
| CRMS32A/DHL-1 | 0.08 | 8.63*** | 2.30 | 0.10 | 0.07 | -0.41 | 41.48* | 0.18** | 41.63* | 4.98 | -2.85*** | -8.22*** |
| APMS6A/DHL-1 | -5.16*** | -12.7*** | -7.46*** | -0.51 | -3.76** | -0.18 | -3.12 | -6.95 | -10.08 | 12.55*** | 0.55 | -5.92 |
| IR58025A/DHL-2 | -17.2*** | 6.44*** | -8.70*** | -0.06 | 1.17 | -0.43 | -5.67 | -4.82 | -10.49 | 3.79* | -0.47 | -9.87*** |
| CRMS32A/DHL-2 | 12.7*** | 3.70*** | -0.69 | 0.10 | -3.17** | 1.91 | 14.32 | 9.02 | 23.35 | -1.49 | 0.98 | 12.68*** |
| APMS6A/DHL-2 | 4.52*** | -10.1*** | 8.70*** | -0.03 | 1.99 | -1.47 | -8.64 | -4.2 | -12.85 | -2.3** | -0.5 | -2.81*** |
| IR58025A/RIL-1 | 0.41* | -4.60* | -1.31 | 0.12 | -2.26* | -0.88 | 5.81 | -14.58* | -8.77 | 9.09*** | -1.16 | 10.57*** |
| CRMS32A/RIL-1 | -9.58*** | -16.2*** | -0.57 | -0.16* | 2.55* | -0.78 | -25.08* | -3.39 | 28.48* | -3.38 | 0.83 | -6.03*** |
| APMS6A/RIL-1 | 9.16*** | 20.81*** | 1.89 | 0.04 | -0.28 | 1.66 | 19.27 | 17.98** | 37.25* | -5.71* | 0.32 | -1.09*** |
| IR58025A/RIL-24 | 11.75*** | -5.96** | 4.16* | -0.1 | -2.60* | 0.71 | 38.21** | 12.63 | 50.84** | -3.59 | -0.65 | -14.3*** |
| CRMS32A/RIL-24 | -3.25*** | 3.86* | -1.02 | 0.03 | 0.55 | -0.71 | -30.72 | -5.81 | -36.53* | -0.11 | 1.02 | 4.53*** |
| APMS6A/RIL-24 | -8.50*** | 2.10 | -3.11 | 0.14 | 2.05 | -0.01 | -7.49 | -6.81 | -14.31 | 3.71 | -0.37 | 11.28*** |

### Trait wise *per se* performance of novel hybrids

As shown in S9-S10 Tables, all novel hybrids were observed to demonstrate heterosis at least for one trait. The hybrid derived from cross, CRMS32A/DHL-1 (Fig 1, S9 Table) indicated *per se* positive heterotic potential for 66.66% of traits under study (i.e., for eight traits).

**Fig 1.**
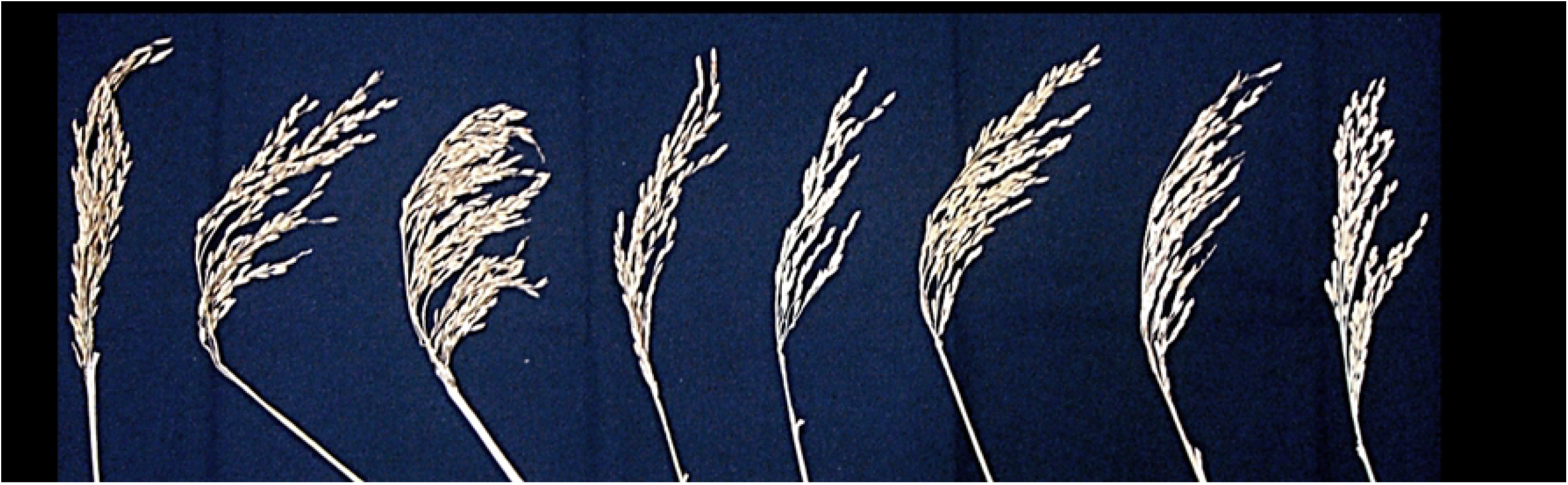
Images of panicles of the novel F_1_hybrid derived from the cross CRMS32A (female parent) and DHL-1 (male parent) along with panicles of the hybrids checks KRH-2, US312, US314, PA6444 and HRI174.

This being an early flowering (DFF, 97 days), it displayed an improvement in six traits viz., YLD, FLW, NPT, TNGPP, NFGPP and SFP. It also recorded a lesser number of unfilled grains than all standard hybrid checks by 37.50%. The percentage of trait improvement when compared with the mean value of all standard checks observed to be as follows: 38.87% (YLD), 25% (FLW), 18.75% (NPT), 82.69% (TNGPP), 115.18% (NFGPP) and 29.72% (SFP). In 75% of the traits which demonstrated heterosis, SCA of the cross was observed to be the causative effect. The hybrid derived from the cross combination CRMS32A/DHL-2 (S10 Table, S1 Fig) was observed to be heterotic for six traits. It was observed to surpass all standard hybrid checks for the traits YLD (by 40.53%), FLL (by 7.09%), NPT (by 18.75%), SFP (by 26.44%) and TGW (by 59.27%). Similar to CRMS32A/DHL-1, this hybrid was also observed to have less number of unfilled grains and exceeded the values of all standard checks by 37.50%. Moreover, 33.33% of traits (i.e., four traits) that showed an improvement were observed to have a higher prevalence of crosses SCA.

The hybrid derived from the cross, APMS6A/DHL-2 (S10 Table, S2 Fig), was an early flowering hybrid (DFF, 90 days i.e. four days earlier than the mean DFF value of all standard checks) along with improvement with respect to the traits FLL (58.78%), NPT (27.77%) and had lesser percentage of unfilled grains as compared to all standard checks. Equal preponderance of GCA-SCA effects were observed with respect to contribution towards improvement of the traits in this hybrid. The hybrid derived from the cross IR58025A/DHL-2 (S10 Table, S3 Fig) was an early flowering hybrid (DFF, 81 days i.e., earlier by 13 days in comparison with the mean value of standard checks) and demonstrated improvement with respect to the traits viz., NPT (by 18.75%), PL (by 16.36%), SFP (by 26.44%) and TGW (14.39%). Higher preponderance of GCA of parents was contributing towards the improved traits. The fifth hybrid derived from the cross combination IR58025A/RIL-24 was observed to be positively heterotic than the mean values of all standard checks for four traits namely YLD (by 51.54%), FLL (by 17.69%), TNGPP (by 50.71%) and NFGPP (by 56.72%). SCA of the cross combination was the causative effect for heterosis in three traits namely FLL, TNGPP and NFGPP. The details of *per se* performance of other novel hybrids are presented in S10 Table and S4-S8 Fig.

As shown in S11 Table, *per se* performance of cross combinations was assessed in view of SCA effect contributions in hybrid improvement. For YLD trait, it was observed that the highest yielding novel hybrid derived from the cross IR58025A/RIL-24 (50.23 g/plant) was observed with high GCA effect (6.13, *P*< 0.001). The GCA nature of both the female parent was observed to be high than its male counter-part. The lowest yielding hybrid (APMS6A/DHL-2, 16.17 g/plant) was observed with negative SCA effect (−2.81) wherein the nature of GCA of parents was observed to be low. For TNGPP trait, the hybrid, IR58025A/RIL-24, with highest number of total number of grains per panicle (i.e., 211) was observed with highest SCA effect value of 50.84, *P*<0.001. Both the parents demonstrated high GCA effects. Also, the hybrid (APMS6A/DHL-2) recorded with lowest number of TNGPP was observed to have a negative SCA effect where both the parents showed low GCA. A similar kind of trend was observed for traits namely FLL, FLW, NFGPP, NUFGPP and SFP.

## Discussion

The primary objective of heterosis breeding programs in crops is to identify best performing hybrids for commercialization along with identifying genetically diverse and better parental lines which could be utilized for the development of promising hybrids in future crosses [32]. Selection of best performing hybrids for yield and other desired characters relies on undertaking trials in multiple environments followed by a rigorous statistical analysis [33]. Trials which are based on various mating designs and which are accompanied by statistical analysis not only differentiates between the effect of genetical and environmental factors but also partitions genetic influences into additive and non-additive components [32, 34]. Such statistical analyses are possible only when an in-depth scrutiny of the relationship between combining ability of parental lines and *per se* heterosis of the novel hybrids is undertaken[35]. As opined by [36], combining ability is an estimate of the genotypic values of lines and testers in relation with their offsprings performance using a definite mating design, which is generally assessed through progeny testing. When the novel progeny demonstrates heterosis, the parental lines are considered to have a good general combining ability. Apart from the general combining ability (GCA) which is statistically main effect in nature, specific combining ability (SCA) is also interactive in nature [37]. Both GCA and SCA play a crucial role in novel hybrid development by influencing the evaluation of inbred lines [38]. Moreover, manifestation of GCA is owed to genetic activities among the loci which are largely additive effect in nature coupled with additive × additive interactions [39] whereas SCA embodies when the genetic loci are under non-additive effects namely dominance and epistatic variances. If epistasis predominates the genetic loci, interactive components namely additive×dominance and dominance×dominance interactions come into play [33]. Considering these points, the present study was undertaken with an objective of assessingthe heterotic potential and combining ability of immortal restorer lines derived from an elite rice hybrid, KRH-2 and identification of promising hybrids with a potential for commercialization.

Evaluation of novel hybrids produced from the test crosses made between popular CMS line, IR58025A and 40 immortal restorers (16 selected DHLs and 24 selected RILs)derived from an elite hybrid KRH-2 demonstrated that four hybrids from restorers namely RIL-1, RIL-24, DHL-1 and DHL-2 out-yielded KRH-2 for YLD trait. Molecular analysis of these four restorers with fertility restoration specific markers for *Rf3* and *Rf4* loci showed the presence of both the genes in them. Also, the fertility restoration analysis of test crosses validated the results of markers analysis and indicated that the DHLs/RILs have complete fertility restoration ability. Our results are in accordance with the reports of [25] and [40], who concluded that that fertility restoration trait is largely controlled by *Rf4* and *Rf3* genes and *Rf4* has a major effect on the trait.

The CV % of the traits DFF, PH, FLL, FLW, PL, SFP and TGW (of parents, novel hybrids and standard hybrid checks) were observed to be below 20%, which indicates that the experiments were, conducted properly [41]. Few traits viz., NPT, TNGPP, NFGPP and YLD recorded higher CV% of > 20%, demonstrating higher genotypic variation between the parental lines, their hybrids and standard checks [42]. Higher phenotypic variance (Vp) and phenotypic coefficient variance (PCV) than their equivalent genotypic variance (Vg) and genotypic coefficient variance (GCV) for all the characters was observed in this study, demonstrating a significant influence of environment on all the traits [43-45]. Though the environmental influence was observed to be higher than genetic preponderance, it was interesting to note that the magnitude of difference between the two kinds of influences was low for all the traits [42]. Therefore, for future breeding programs, selection of genotypes could be based on these traits as pointed out by [44, 46]. Further, [29, 47] indicated that heritability plays an important role in studying quantitative characters by indicating genetically, the reliability of a phenotype for enhancing its breeding value. High heritability estimates for all the traits except FLW demonstrate the traits’ higher response during selection. These results are in accordance with the reports of [43, 47, 48]. As opined by [49], the magnitude of genotypic trait improvement in novel population vis-à-vis parental population at certain selection intensity less than one selection cycle is revealed through an assessment of genetic advance (GA). In our study the GA value for YLD trait was observed to be 24.48. This indicates that if better performing hybrids identified in this study are used as restorers in future crossing programs, then they can significantly contribute towards development of better novel hybrids [42]. Further, as pointed out by [50], prediction of genetic gain under selection is more reliable when high heritability (*H*^*2*^) estimates are combined with high genetic advance as percent of the mean (GAM). It is because GAM helps in understanding the type of gene action for various polygenic traits and therefore combined estimates of high *H*^*2*^-high GAM is an important selection parameter. In this study, the traits DFF, PH, FLL, NPT, TNGPP, NFGPP, SFP and YLD demonstrated high *H*^*2*^-high GAM and high *H*^*2*^ with moderate GAM noticed for PL trait. These observations imply the predominance of additive gene action and selection of the above mentioned traits could be possible if done in early generations [49, 51]. However, moderate *H*^*2*^ with moderate GAM was observed with FLW trait indicating non-additive gene action implying that the improvement of FLW trait is only possible through deployment of recurrent selection method [42, 51].

Correlation analysis is used as an important tool for indirect selection as it aids the plant breeder in getting a better understanding of the various traits, which may influence yield. As pointed out by [52], improvement in the desired character (such as grain yield) on the basis of its allied traits selection would be evident in successive generations of segregating population. Keeping these points in view, in this study, the correlation analysis between the various component traits of yield were assessed in the different rice hybrids and the analysis revealed a strong and positive correlation between YLD and its important allied parameters namely NPT, PL, FLL, TNGPP, SFP, FLW and NFGPP at both 1% and 5% levels of significance [53, 54]. This observation indicates that selection of these traits in parental lines will be helpful for development of hybrids with better yield heterosis. A negative correlation was observed between YLD and PH trait in this study and this is in accordance with the earlier reports [55,56]. A positive but insignificant correlation was observed between PH and SFP which is in accordance with the study of [57-59]. Negative correlation between DFF and PH, PT, NFGPP, TGW, YLD was observed in our study. These results are in accordance with [60] (for DFF and YLD), with [61] (for DFF and NPT), with [42] (for DFF and NFGPP, TGW) and is not in accordance with [2, 61] (for DFF and PH).

All the novel hybrids were observed to be positively heterotic for any of the three categories of heterosis, for at least one important trait related to productivity. However, none of the hybrids showed positive heterosis for all the traits under study and they out yielded the parents and standard hybrid checks for YLD and its allied trait components in various degrees except for PH. For five traits namely, YLD, FLL, FLW, TNGPP and NFGPP, average standard heterosis (SH) for all cross combinations was the highest and the values were in the range of 1.31% (FLW) to 189.01% (YLD). For six traits namely, NPT, PL, NUFGPP, SFP, TGW and DFF, average mid-parent heterosis (MPH) value was highest among the three categories for all cross combinations and it was in the range of 7.52% (NUFGPP) to 46.08% (NPT). The trait PH demonstrated negative heterosis in the novel hybrids. Different values for variance with respect to mid-parent heterosis (MPH), better parent heterosis (BPH) and standard heterosis (SH) was observed among the novel hybrids. These observations are in accordance with earlier observations those previously reported by [3, 62, 63].

Estimates of GCA have been used in various crop breeding systems as an index of breeding value of a particular genotype [38]. A higher value for GCA demonstrates the predominance of parental mean over than the general mean. As per [64], high GCA values are not only indicative of the flow of useful genes from parents to offspring at a higher rate but are also an indicator of additive gene action in play. Therefore, from the breeding point of view, an estimate of higher GCA also indicates higher heritability and reduced environmental effects [33] and higher gene interactions [34, 65]. As observed by [36, 66], a parent that is good in *per se* performance, when used in hybridization may not produce a superior hybrid. In such cases, a superior hybrid can be produced when there would be a proper selection of other parent for hybridization with a poor GCA [67]. In the present study for determining the best combiner among the lines and testers, positive and significant values were considered for all the traits except DFF, PH and NUFGPP. Among the CMS lines under study, IR58025A demonstrated best combining ability for traits viz., PL (2.11***), SFP (6.16***) and YLD (6.13***). CRMS32A was observed to show positive and significant GCA for traits namely NFGPP (17.76***) and TNGPP (16.2*). APMS6A was observed to show best combining ability for the traits namely DFF (−6.83***), FLL (3.01**) and NPT (6.22***). Since, IR58025A showed higher GCA values for YLD trait and for its two crucial trait components; it was considered to be the best combiner among the lines. Among the testers, DHL-1 was observed to have positive and significant GCA values for the traits namely FLW (0.18***), NFGPP (14.18*), SFP (4.21***) and YLD (4.61***). Tester DHL-2 showed positive and significant GCA values for traits NPT (1.67***) and TGW (2.10***) and desirable negative GCA values for the traits DFF (−6.80***) and PH (−10***). RIL-24 was observed to have the higher GCA values for traits PL (1.17*), NFGPP (20.07**) and TNGPP (21.43**). Lastly, RIL-1 showed positive and significant GCA value for trait PL (1.76***). Therefore, DHL-1 was identified to be the best combiner among the testers.These observations are in accordance with reports of [3, 68, 69], wherein no single parent was identified to have a good GCA for all the traits in their studies. [2]demonstrated that the most promising and high yielding hybrids in their study were produced from two of the parents which showed the best GCA for grain yield. In our study, the most promising hybrid (CRMS32A/DHL-1) which showed heterosis for eight traits, an equal predominance of SCA and GCA effects was observed which translated into higher percentages of heterosis than all the standard varietal and hybrid checks. Highest yielding hybrid (IR58025A/RIL-24) was produced from those parents who showed the best and poor GCA for YLD trait. Therefore, our observations are partly in accordance with the report of [2].

If the potential of a parental line to combine well in a particular cross combination is observed then they are supposed to have a good SCA [33]. Based on the GCA of parents, various types of gene actions have been reported for the manifestation of SCA (in cross combinations) among the various crop systems. As observed by [70, 71], both the parents with a good GCA may produce a high SCA effect attributing to additive × additive gene action whereas desirable additive effects of a good combiner and favorable epistatic interactive effects of poor combiner parent comes into play when good and poor GCA parents results in a high SCA effect [72]. Lastly, both the parents having low GCA effects, if observed to bring about high SCA effect, it may be due to over dominance, non-allelic gene interaction i.e., of dominance × dominance type [73]. In our study for two traits namely FLW and PL, no significant SCA effects were observed, indicating that the trait values are in the range of parental averages [2].

The SCA estimates of cross combinations and GCA effects of the parents are described in S9-S10 Tables. For the YLD trait, the highest yielding hybrid (50.23 g /plant) derived from IR58025A/RIL-24 resulted from good-by-poor general combiners, which implies that additive × additive gene action, was the causative effect of heterosis in this cross. In the lowest yielding novel hybrid (16.17 g/plant) derived from the cross combination, APMS6A/DHL-2, both the parents were poor combiners for yield. For NPT trait, four cross combinations, APMS6A/DHL-2, APMS6A/DHL-1, IR58025A/DHL-2 and CRMS32A/RIL-24 produced the hybrid with highest number of productive tillers (i.e.,18). Higher predominance of GCA effect on the above mentioned four crosses, additive × additive gene action of parents brought about heterosis in the NPT trait. Though the hybrid from cross combination APMS6A/DHL-2 had more number of NPT, the least number of TNGPP and NFGPP might be attributed to its least yield. The hybrid produced from the cross combination, IR58025A/RIL-1, was observed to have a negative SCA effect with least number of productive tillers, this demonstrates an occurrence of bad combination of alleles from the parents responsible for negative SCA effect in the hybrid [2]. For TNGPP trait, the hybrid IR58025A/RIL-24 recorded highest number of total grains per panicle (i.e., 211). An exceptionally high SCA value was observed in this hybrid, which validated the SCA’s predominance. The nature of GCA effect of one of the parents was high which indicated towards additive × additive gene action for SCA’s prevalence. The least number of total grains per panicle was observed in the hybrid APMS6A/DHL-2. The nature of GCA of parents was poor-by-poor that might have resulted in lesser trait expression. Dominance × dominance gene interaction promoted the SCA’s dominance on the hybrid but the combination of bad parental alleles contributed for a negative SCA value. For TGW trait, the hybrid IR58025A/DHL-2 with the highest TGW (23.20 g) was observed to have a higher GCA effect than SCA as the nature of GCA effect of one of the parents was high. For the cross combination, CRMS32A/DHL-1 the lowest TGW value of 15.23 g might be due to unfavorable allelic combination from parents resulted in negative SCA value. These results are partly in accordance with [2, 3, 62, 63], who reported that in the crosses analyzed in their studies, despite both the parents being good general combiners, they could not produce hybrids with positive SCA values. In our study, in those cross combinations with higher and positive SCA values (due to high GCA values of parents), good combination of alleles from the parents might have resulted in positive SCA [2]. Also, those cross combinations that showed negative SCA effect value in our study, witnessed low to average GCA parental effects which is not in accordance with [68, 69]. The predicted genetic mechanisms of the SCA delineated in this study were in accordance with the report of [33].

In our study, the SCA variance was observed to be higher than the GCA variance. Higher values of dominance genetic variance (δ^2^D) were observed for all the traits except FLW and PL. With respect to traits viz., TNGPP, NUFGPP, PH, YLD, DFF and FLL,these were under partial dominance, whereas the traits FLW, NPT, PL, NFGPP, SFP and TGW were a consequence of dominance and also the GCA variance ratio for all the traits was below 1. All these observations indicate a predominance of non-additive gene action which is partly in accordance with the report of [2], wherein few traits were observed to be under preponderance of additive gene action. Also, the narrow sense heritability (*h*^*2*^) values were greater than 10 for all the traits with values being lower than broad sense heritability (*H*^*2*^). It is surprising to note that trait PL recorded lower values of both *h*^*2*^ and *H*^*2*^. This might be due to lower phenotypic variation among the novel hybrids for this trait as explained by [2]. Our observation of non-additive gene action underlying the genetic variance of all the traits under study are in accordance with earlier observations by [74, 75], thus highlighting the importance of non-additive gene action in heterosis breeding.

Improvement in the quantitative traits is dependent on factors such as proportionate contribution of GCA and SCA to the crosses along with the type of breeding method to be used for augmenting the trait value and whether early or late generations of breeding material are to be used for the same. These factors help a plant breeder to make crucial decisions in the course of plant breeding. As demonstrated by [76, 77] in other crops such as maize, if the GCA variances exceed the SCA variances, the prediction of GCA effects is based on an assessment of genotypes during early generations which aids in identification of promising hybrids. This is because if a line with good GCA is identified during early generations, this early selection aids in the transfer of favorable and heritable genetic material from parents to offsprings [78]. Early testing of such promising lines with high GCA values not only aids in improvement of parental lines but also reduces the cost and time for its development during breeding programs. When high SCA effect indicates the preponderance of non-additive genetic variance, selection of such lines should be undertaken in later generations when such variances are known to get fixed in homozygous lines [79, 80].

As suggested by [81], a standard heterosis percentage of 20%-30% in a hybrid is considered optimum to mitigate the higher seed cost and bring profit to farmers in self-pollinated crops such as rice. In our study, the promising heterotic combinations (for YLD trait) were assessed for standard heterosis percentages, for trait *per se* performance and GCA-SCA effects. In our investigation, 58.33% (i.e., seven) of the novel cross combinations had heterosis percentages more than 20% for YLD trait when analyzed for all categories of yield heterosis. Among them, two hybrids namely IR58025A/RIL-24 (50.23 g/plant), CRMS32/RIL-24 (43.90 g/plant) were observed to have standard heterosis of more than 50% when assessed with five standard checks. Also, among the seven novel crosses which were positively heterotic for all categories of YLD trait heterosis, three crosses (42.85%), namely CRMS32A/RIL-24, APMS6A/RIL-24 and CRMS32A/DHL-2 demonstrated high SCA effect. It was interesting to note that, these three cross combinations demonstrated high SCA effects and low GCA value for both the parents. Higher SCA values may be due to over dominance, non-allelic gene interaction i.e., of dominance × dominance type as delineated by [73]. Therefore, as suggested by [3], in order to produce heterotic F_1_s, selection of parents with diverse GCA effects i.e. high-low and low-low may be undertaken to realize better levels of high heterosis. For TNGPP trait, IR58025A/RIL-24 cross observed to demonstrate standard heterosis more than 50% with desirable SCA effects. Therefore, the seven cross combinations, viz., IR58025A/RIL-24, CRMS32A/RIL-24, APMS6A/RIL-24, CRMS32A/DHL-1, APMS6A/DHL-1, IR58025A/DHL-2 and CRMS32A/DHL-2 which demonstrated positive heterosis for all the categories of YLD heterosis could be considered for commercial exploitation. The extra-ordinarily higher levels of heterosis (as high as 189.01% of standard heterosis in IR58025A/RIL-24as compared to check US314) observed in this study were previously reported by [82, 83]. The identified promising novel hybrids in this study needs to be evaluated in multi-location trials to assess their consistency with respect to yield heterosis and after validation; they can be released for cultivation.

## Conclusion

In the present study, combining abilities of three popular CMS lines (IR58025A, CRMS32A and APMS6A)and four novel restorers (DHL-1, DHL-2, RIL-1 and RIL-24) through line × tester analyses were assessed for their utility in hybrid breeding. CMS line IR58025A was identified as best combiner as it showed positive significant values for total grain yield per plant, panicle length and spikelet fertility. Distinct parents that produced heterotic hybrids and possessing varying levels of SCA effects were identified and they could be utilized for heterosis breeding. Seven cross combinations (IR58025A/RIL-24, CRMS32A/RIL-24, APMS6A/RIL-24, CRMS32A/DHL-1, APMS6A/DHL-1, IR58025A/DHL-2 and CRMS32A/DHL-2) which demonstrated positive heterosis for all the categories of yield heterosis were identified for commercialization. These hybrids could be considered for large-scale cultivation after their validation in multi-location trials.

## Acknowledgement

Swapnil is grateful to DST INSPIRE Fellowship Division, New Delhi for providing the financial assistance (Grant # DST/INSPIRE Fellowship/2013/1146), Department of Biotechnology (DBT), Govt. of India and to the Director, ICAR-IIRR for providing the infrastructural facilities for carrying out this research work.

## Supporting information

S1 Fig. Images of panicles of the novel F_1_ hybrid derived from the cross CRMS32A (female parent) and DHL-2 (male parent) along with panicles of hybrid checks, KRH-2, US312, US314, PA6444 and HRI174. CRMS32A/DHL-2 was used as heading of the image. (TIF).

S2 Fig. Images of panicles of the novel F_1_ hybrid derived from the cross APMS6A (female parent) and DHL-2 (male parent) along with panicles of hybrid checks, KRH-2, US312, US314, PA6444 and HRI174. APMS6A/DHL-2 was used as heading of the image. (TIF).

S3 Fig. Images of panicles of the novel F_1_ hybrid derived from the cross IR58025A (female parent) and DHL-2 (male parent) along with panicles of hybrid checks, KRH-2, US312, US314, PA6444 and HRI174. IR58025A/DHL-2 was used as heading of the image. (TIF).

S4 Fig. Images of panicles of the novel F_1_ hybrid derived from the cross CRMS32A (female parent) and RIL-24 (male parent) along with panicles of hybrid checks, KRH-2, US312, US314, PA6444 and HRI174. CRMS32A/RIL-24 was used as heading of the image. (TIF).

S5 Fig. Images of panicles of the novel F_1_ hybrid derived from the cross APMS6A (female parent) and RIL-24 (male parent) along with panicles of hybrid checks, KRH-2, US312, US314, PA6444 and HRI174. APMS6A/RIL-24 was used as heading of the image. (TIF).

S6 Fig. Images of panicles of the novel F_1_ hybrid derived from the cross CRMS32A (female parent) and RIL-1 (male parent) along with panicles of hybrid checks, KRH-2, US312, US314, PA6444 and HRI174. CRMS32A/RIL-1 was used as heading of the image. (TIF).

S7 Fig. Images of panicles of the novel F_1_ hybrid derived from the cross APMS6A (female parent) and DHL-1 (male parent) along with panicles of hybrid checks, KRH-2, US312, US314, PA6444 and HRI174. APMS6A/DHL-1 was used as heading of the image. (TIF).

S8 Fig. Images of panicles of the novel F_1_ hybrid derived from the cross APMS6A (female parent) and RIL-1 (male parent) along with panicles of hybrid checks, KRH-2, US312, US314, PA6444 and HRI174. APMS6A/RIL-1 was used as heading of the image. (TIF).

S1 Table: Estimation of standard YLD heterosis in novel F_1_ hybrids derived from test crosses of selected 16 DHLs with IR58025A assessed in the dry season of 2018-2019 (DOCX).

S2 Table: Estimation of standard YLD heterosis in novel F_1_ hybrids derived from test crosses of selected 24 RILs (12 high and 12 low yielding) with IR58025A assessed in the dry season of 2018-2019 (DOCX). Footnote common to S1 Table, S2 Table: *Rf3/Rf4*-fertility restoration major loci, DHL-doubled haploid line, RIL-recombinant inbred line, AKD-Akshayadhan, VRD-Varadhan, KRH-2-Karnataka Rice Hybrid-2. Gist of S1-S2 Tables: Promising cross combinations are indicated in bold. Four crosses namely IR58025A/DHL-1, IR58025A/DHL-2, IR58025A/RIL-1 and IR58025A/RIL-24 demonstrated positive standard total grain yield per plant (YLD) heterosis when Akshayadhan (AKD) and Varadhan (VRD) were used as standard varietal checks and KRH-2 as standard hybrid check. Molecular assessment of four donors, DHL-1, DHL-2, RIL-1 and RIL-24 with functional markers specific for *Rf3-Rf4* loci showed the presence of both the loci. Fertility restoration potential of these donors indicated them to be complete restorers. Therefore, these four donors were chosen for test cross with popular CMS lines, IR58025A, CRMS32A and APMS6A in the wet season 2018 (DOCX).

S3 Table: Genetic variability estimation for 12 agro-morphological traits in novel hybrids derived from the cross between IR58025A, CRMS32A, APMS6A and DHL-1, DHL-2, RIL-1 and RIL-24. Footnote of the S3 Table: Vg-Genotypic variance and Vp-phenotypic variance, GCV-genotypic coefficient of variance and PCV-phenotypic coefficient of variance, H2, broad-sense heritability, GA-genetic advance and GAM-genetic advance as percent of mean, DFF-Days to fifty percent flowering; PH-Plant Height (cm), FLL-Flag leaf length (cm); FLW-Flag Leaf Width (cm); NPT-Number of Productive Tillers; PL-Panicle Length (cm); TNGPP-Total number of grains per panicle; NFGPP-Number of filled grains per panicle; NUFGPP-Number of unfilled grains per panicle; SFP-Spikelet fertility in percentage; TGW-Test (1,000) grain weight (g); YLD-Total grain yield per plant (g) (DOCX).

S4 Table: Correlation among the agro-morphological traits studied in novel hybrids derived from the cross between IR58025A, CRMS32A, APMS6A and DHL-1, DHL-2, RIL-1 and RIL-24. Footnote of S4 Table: DFF-Days to fifty percent flowering; PH-Plant Height (cm), FLL-Flag Leaf Length (cm); FLW-Flag Leaf width (cm); NPT-Number of Productive Tillers; PL-Panicle Length (cm); TNGPP-Total Number of Grains Per Panicle; NFGPP-Number of Filled Grains per Panicle; NUFGPP-Number of Unfilled Grains per Panicle; SFP-Spikelet Fertility in Percentage; TGW-Test (1,000) Grain Weight (g); YLD-Total grain Yield per plant (g); *\*P < 0*.*05, **P < 0*.*01* (DOCX)

S5 Table: Genetic variances and heritability estimation for total grain yield per plant (YLD) and its allied traits in novel hybrids derived from the cross between IR58025A, CRMS32A, APMS6A and DHL-1, DHL-2, RIL-1 and RIL-24. Footnote of S5 Table: σ^2^gca, variance due to general combining ability (GCA); σ^2^sca, variance due to specific combining ability (SCA); σ^2^gca / σ^2^sca, GCA variance ratio; δ^2^A, additive genetic variance; δ^2^D, dominance genetic variance; (δ^2^D/ δ2^A^)1/2, degree of dominance; *H*^*2*^, broad-sense heritability; *h*^*2*^, narrow-sense heritability; DFF-Days to fifty percent flowering; PH-plant height (cm), FLL-Flag leaf length (cm); FLW-Flag leaf width (cm); NPT-Number of productive tillers; PL-panicle length (cm); NFGPP-Number of filled grains per panicle; NUFGPP-Number of unfilled grains per panicle TNGPP-Total number of grains per panicle; SFP-Spikelet fertility in percentage; TGW-test (1,000) grain weight (g); YLD-Total grain yield per plant (g) (DOCX).

S6 Table: ANOVA for agro-morphological traits in novel hybrids derived from the cross between IR58025A, CRMS32A, APMS6A and DHL-1, DHL-2, RIL-1 and RIL-24. Footnote of S6 Table: *df*-degrees of freedom, SS-Sum of Squares, MSS-Mean Sum of Squares, F value-F values statistic, prob-probability value, DFF-Days to fifty percent flowering; PH-Plant Height (cm), FLL-Flag leaf length (cm); FLW-Flag Leaf Width (cm); NPT-Number of Productive Tillers; PL-Panicle Length (cm); TNGPP-Total number of grains per panicle; NFGPP-Number of filled grains per panicle; NUFGPP-Number of unfilled grains per panicle; SFP-Spikelet fertility in percentage; TGW-Test (1,000) grain weight (g); YLD-Total grain yield per plant (g) (DOCX).

S7 Table: Estimation of standard heterosis (SH), mid-parent heterosis (MPH) and better parent heterosis (BPH) for total grain yield (YLD) of the novel hybrids derived from the cross between IR58025A, CRMS32A, APMS6A and DHL-1, DHL-2, RIL-1 and RIL-24. Footnote of S7 Table: SH(HRI174) (%)-Standard heterosis in percentage when HRI174 is used as varietal check; SH(PA6444) (%)-Standard heterosis in percentage when PA6444 is used as varietal check; SH(US314)(%)-Standard heterosis in percentage when US314 is used as varietal check; SH (US312)-Standard heterosis in percentage when US312 is used as varietal check (%); SH(KRH-2)(%)-Standard heterosis in percentage when US314 is used as varietal check; MPH (%)-Mid parent heterosis; BPH (%)-Better parent heterosis, RIL-1-highest yielding RIL, RIL-24-low yielding RIL, DHL-1-Highest yielding doubled haploid line, DHL-2, Second highest yielding doubled haploid line; *\*P< 0*.*05, **P< 0*.*01* (DOCX).

S8 Table: Estimation of standard heterosis, mid-parent heterosis and better parent heterosis for yield related components in novel hybrids. Footnote of S8 Table: SH(HRI174) (%)-Standard heterosis in percentage when HRI174 is used as varietal check; SH (PA6444) (%)-Standard heterosis in percentage when PA6444 is used as varietal check; SH (US314)(%)-Standard heterosis in percentage when US314 is used as varietal check; SH (US312)-Standard heterosis in percentage when US312 is used as varietal check (%); SH (KRH-2)(%)-Standard heterosis in percentage when US314 is used as varietal check; MPH (%)-Mid parent heterosis; BPH (%)-Better parent heterosis, RIL-1-highest yielding RIL, RIL-24-low yielding RIL, DHL-1-Highest yielding doubled haploid line, DHL-2, Second highest yielding doubled haploid line; *\*P < 0*.*05, **P < 0*.*01* (DOCX).

S9 Table: Trait-wise *per se* performance, SCA-GCA effects in most promising novel hybrid. Foot note of S9 Table: YLD-Total grain yield per plant (g), DFF-Days to fifty percent flowering; PH-Plant Height (cm), FLL-Flag leaf length (cm); FLW-Flag Leaf Width (cm); NPT-Number of Productive Tillers; PL-Panicle Length (cm); NFGPP-Number of filled grains per panicle; NUFGPP-Number of unfilled grains per panicle TNGPP-Total number of grains per panicle; SFP-Spikelet fertility in percentage; TGW-Test (1,000) grain weight (g); *\*P < 0.05; **P < 0.01; ***P < 0.001*; †-early flowering hybrid than HRI174, Ɨ-denotes hybrids heterotic than controls namely HRI174, PA6444, US314, US312, KRH-2, ¶-denotes less number of unfertile grains than standard hybrid checks. Gist of S9 Table: The novel hybrid was observed to be heterotic than the standard checks for eight crucial yield related viz., YLD, DFF (early flowering hybrid), NPT, TNGPP, NFGPP, NUFGPP, SFP% traits due to higher prevalence of specific combining ability (SCA) making it the best performing hybrid (DOCX).

S10 Table: Trait-wise *per se* performance, SCA-GCA effects in novel hybrids. Foot note of S10 Table: YLD-Total grain yield per plant (g), DFF-Days to fifty percent flowering; PH-Plant Height (cm), FLL-Flag leaf length (cm); FLW-Flag Leaf Width (cm); NPT-Number of Productive Tillers; PL-Panicle Length (cm); NFGPP-Number of filled grains per panicle; NUFGPP-Number of unfilled grains per panicle TNGPP-Total number of grains per panicle; SFP-Spikelet fertility in percentage; TGW-Test (1,000) grain weight (g); *\*P < 0*.*05; **P < 0*.*01; ***P < 0*.*001*, SCA-Specific Combining Ability, GCA-General Combining Ability; I-denotes hybrids heterotic than controls namely HRI174, PA6444, US314, US312, KRH-2; ¶-denotes less number of unfertile grains than checks; †-early flowering hybrid than all checks; ƪ-early flowering hybrid than HRI174, KRH-2 only; ‖-early flowering hybrid than donor parent, RIL-24, only; ●-early flowering hybrid than all checks except US312; ʘ-early flowering hybrid than all checks except US314 (DOCX).

S11 Table: *Per se* performance of cross combinations and SCA effects in hybrid improvement. Footnote of S11 Table: *\*P < 0*.*05; **P < 0*.*01; ***P < 0*.*001*. As opined by Gramaje et al. (2020) selection of cross combinations with high SCA does not lead to the direct improvement in self-pollinated crops such as rice but we have demonstrated that crosses with high SCA have considerably improved the traits namely YLD, TNGPP, DFF (early flowering), PH, FLL, FLW, NFGPP, SFP and NUFGPP. Therefore, from our study, it can be concluded that the selection of crosses with high SCA did automatically transfer the SCA effects to better hybrid performance which is contrary to that of Gramaje et al. (2020). Cross combinations in this study with improved traits due to high SCA are ideal for heterosis breeding (DOCX).

